# Insecticide resistance status of *Aedes aegypti* in Bangladesh

**DOI:** 10.1101/2020.07.31.231076

**Authors:** Hasan Mohammad Al-Amin, Fatema Tuj Johora, Seth R. Irish, Muhammad Riadul Haque Hossainey, Lucrecia Vizcaino, Kishor Kumar Paul, Wasif A. Khan, Rashidul Haque, Mohammad Shafiul Alam, Audrey Lenhart

**Author notes:** Contributed equally (Mohammad Shafiul Alam, Audrey Lenhart).

## Abstract

**Background:** Arboviral diseases including dengue and chikungunya are major public health concern in Bangladesh, with unprecedented levels of transmission reported in recent years. The primary approach to control these diseases is control of *Aedes aegypti* using pyrethroid insecticides. Although chemical control is long-practiced, no comprehensive analysis of *Ae. aegypti* susceptibility to insecticides has previously been conducted. This study aimed to determine the insecticide resistance status of *Ae. aegypti* in Bangladesh and investigate the role of detoxification enzymes and altered target site sensitivity as resistance mechanisms.

**Methods:** *Aedes* eggs were collected using ovitraps from five districts across the country and in eight neighborhoods of the capital city Dhaka from August to November 2017. CDC bottle bioassays were conducted for permethrin, deltamethrin, malathion, and bendiocarb using 3-5-day old F_0_-F_2_ non-blood fed female mosquitoes. Biochemical assays were conducted to detect metabolic resistance mechanisms and real-time PCR was performed to determine the frequencies of the knockdown resistance (*kdr*) mutations Gly1016, Cys1534, and Leu410.

**Results:** High levels of resistance to permethrin were detected in all *Ae. aegypti* populations, with mortality ranging from 0 – 14.8% at the diagnostic dose. Substantial resistance continued to be detected against higher (2X) doses of permethrin (5.1 – 44.4% mortality). Susceptibility to deltamethrin and malathion varied between populations while complete susceptibility to bendiocarb was observed in all populations. Significantly higher levels of esterase and oxidase activity were detected in most of the test populations as compared to the susceptible reference Rockefeller strain. A significant association was detected between permethrin resistance and the presence of Gly1016 and Cys1534 homozygotes. The frequency of *kdr* alleles varied across the Dhaka populations, and Leu410 was not detected in any of the tested populations.

**Conclusions:** The detection of widespread pyrethroid resistance and multiple mechanisms highlights the urgency for implementing alternate *Ae. aegypti* control strategies. In addition, implementing routine monitoring of insecticide resistance in *Ae. aegypti* in Bangladesh will lead to a greater understanding of susceptibility trends over space and time, thereby enabling the development of improved control strategies.

## Background

*Aedes* (*Stegomyia*) *aegypti* (Linnaeus, 1762) is an important vector of arboviral diseases, principally dengue, chikungunya, and Zika. These increasingly common arboviral infections cause severe febrile illness and short to long-term physical or cognitive impairments and even death. Dengue is the most globally prevalent and rapidly spreading arboviral disease, with an estimated 390 million annual infections and 3.9 billion people at risk [1]. Chikungunya is also increasingly prevalent, and the prolonged pain and rheumatism resulting from infection can result in long-term physical problems and impaired daily life [2, 3]. Recently, Zika caused a major global pandemic in 2015-2016, leading to congenital malformations, Guillain-Barre syndrome, and other severe neurological complications [4].

The burden of arboviral diseases in Bangladesh is not well documented. The first major outbreak of dengue took place during the 2000 monsoon, and caused5,521 officially reported cases with 93 deaths [5]. Since then, thousands of infections are reported each year although these numbers represent a fraction of the actual burden since only admitted cases in some selected hospitals are officially reported [6]. Recent estimates suggest that 40 million people have been infected nationally with an average of 2.4 million infections annually. Cases are mostly concentrated in the capital city Dhaka, where the seropositivity ranges from 36 to 85% [7]. In 2019, Bangladesh experienced its largest outbreak with 101,354 confirmed cases and 164 deaths [8]. Since 2008, sporadic infections with chikungunya virus have been reported across Bangladesh, with the largest outbreak occurring in 2017 which infected hundreds of thousands of inhabitants of Dhaka [9]. Zika virus transmission has not been widely reported, with only a single confirmed case in 2016 in a 67-year old man from Chittagong who had not traveled outside of Bangladesh. Although a few additional Zika virus infections were detected by antibody tests, there is no further evidence of Zika in Bangladesh [10, 11].

*Aedes aegypti* is the principal vector of dengue, Zika, and chikungunya. It is highly abundant throughout Bangladesh, especially in Dhaka [7]. In 2018, the Breteau Index (BI; the number of *Aedes*-positive containers per 100 houses inspected) was greater than 100 in some parts of Dhaka [12]. Recent studies in Dhaka confirmed that plastic containers (plastic drums, buckets, plastic bags, bottles, and disposable cups) and discarded vehicle and construction materials (tires, battery shells, and cement mixers) are key containers for *Aedes* production. These are typical of the domestic and industrial detritus that encourage the proliferation of *Ae. aegypti* across the globe. High *Aedes* abundance in Dhaka is also strongly associated with favorable climatic factors including rainfall, temperature, and humidity [13].

In the absence of effective therapeutic drugs and vaccines, *Ae. aegypti* control is presently the only approach for preventing and controlling the transmission of most *Aedes*-borne arboviruses. *Aedes aegypti* control strategies rely heavily on the application of a limited number of chemical insecticides approved for public health use, principally pyrethroids, organochlorines, organophosphates, and carbamates [14]. Of these, pyrethroid insecticides such as deltamethrin, cypermethrin, and permethrin are commonly used because of their low toxicity to mammals and their high efficacy against vectors. However, resistance to many insecticides has emerged in *Ae. aegypti* across the globe and is a serious threat to control programs [15, 16, 17, 18, 19].

Resistance to insecticides is a dynamic evolutionary process, driven by insecticide selection pressures [20]. Resistance can be caused by physiological changes including 1) changes to the mosquito cuticle so insecticides cannot penetrate, 2) increased activity of insecticide detoxification enzymes, and/or 3) structural modifications at the target site of the insecticide or by behavioral adaptations like insecticide avoidance [21].

Target site alterations resulting in resistance to pyrethroids and dichlorodiphenyltrichloroethane (DDT) are often caused by mutations in the voltage-gated sodium channel (VGSC) transmembrane protein and are broadly referred to as ‘knockdown resistance’ (*kdr*) mutations. There are several point mutations on the VGSC gene known to confer *kdr*-type insecticide resistance in *Ae. aegypti*, most notably at positions 410, 1016 and 1534 [22, 23]. Increased enzyme activity resulting in metabolic resistance typically involves any of the three main groups of detoxification enzymes: carboxylesterases, mixed-function oxidases (MFOs), and glutathione S-transferases (GSTs) [24]. Understanding the mechanisms of resistance and their specificity amongst insecticides is important to devising strategies to mitigate and manage insecticide resistance when it is detected.

Although there is a recognized increase of *Aedes*-borne arboviruses in Bangladesh over the last 20 years, little or no organized use of insecticides against *Ae. aegypti* has occurred. Regular control activities are mostly carried out only in Dhaka, targeting the nuisance biting *Culex quinquefasciatus* and *Aedes* by thermal fogging with a combination of pyrethroid insecticides including permethrin, prallethrin, and tetramethrin/bioallethrin. Rising *Aedes*-borne viral diseases indicate little impact of these insecticides being used. Development of resistance against commonly used insecticides in local *Aedes* population may contribute to the failure of the vector control strategy. Occasional source reduction is also carried out by community engagement by both government and private initiatives. However, gaining access to all premises and achieving sufficient coverage of myriad oviposition sites in densely populated cities like Dhaka is a huge challenge [25]. There are also structural challenges to control activities related to management, evaluation, and budget [26, 27]. The insecticide resistance status of *Ae. aegypti* has not previously been comprehensively assessed in Bangladesh. The purpose of this study was to assess the insecticide resistance status and resistance mechanisms of key *Ae. aegypti* populations in Bangladesh to better guide future insecticide choices for vector control.

## Methods

### Study sites

Mosquitoes were collected from five districts throughout Bangladesh. Of these, the capital city, Dhaka and Chittagong are the high-transmission settings and Rajshahi and Chapai Nawabganj are low-transmission settings [9, 28, 29, 30, 31, 32] (Fig. 1). The other district, Bandarban, was selected as it is endemic for malaria and deltamethrin based long-lasting insecticidal nets (LLINs) are regularly distributed with seasonal sporadic indoor residual spraying (IRS) [33, 34]. Since the majority of *Aedes*-borne arboviral infections are reported from Dhaka, eight areas within Dhaka City were selected for sampling [35].

**Fig. 1.**
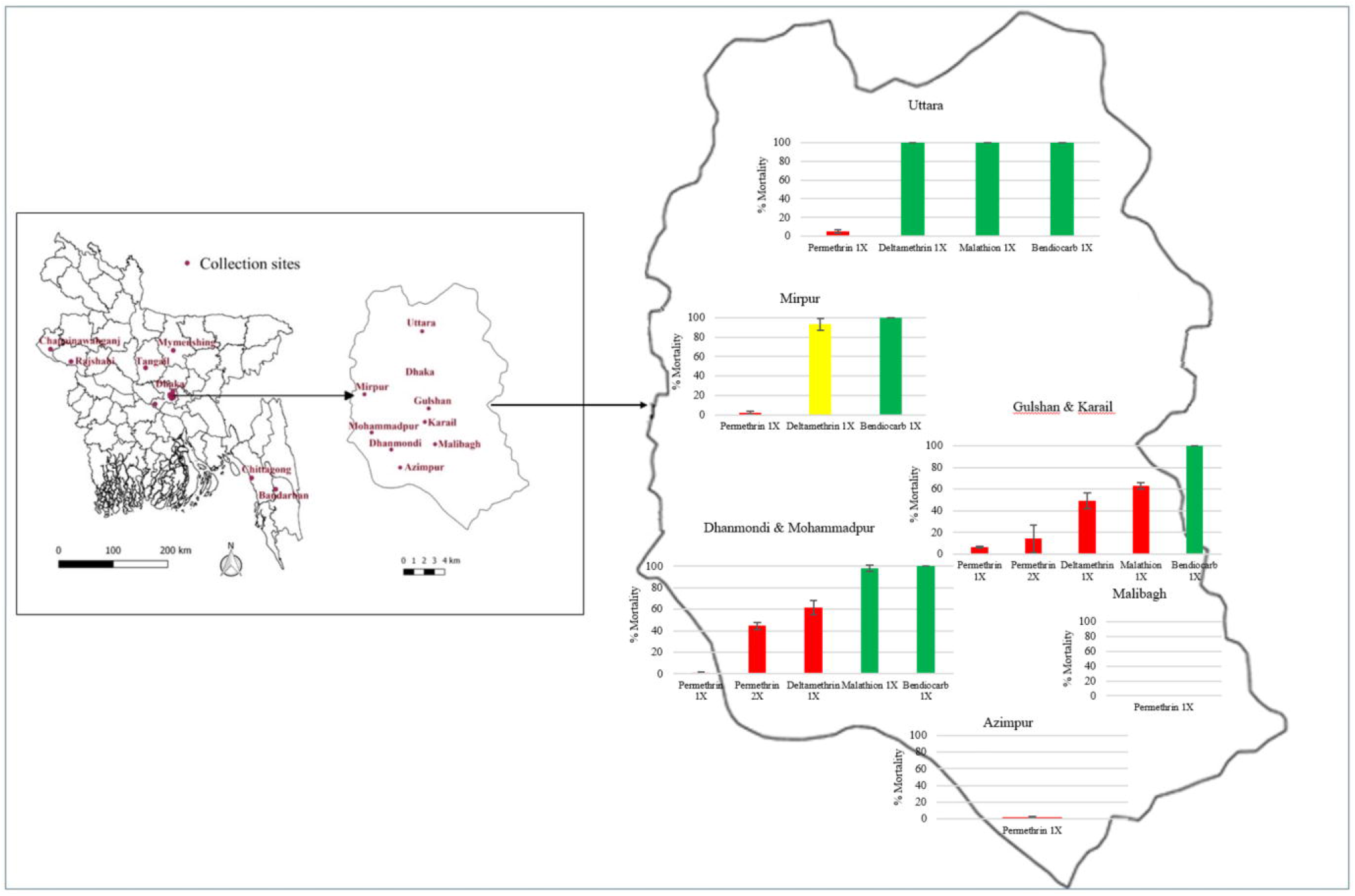
Bioassay results (% Morality at 30 minutes ±95% confidence intervals) for female *Ae. aegypti* in Dhaka. Red columns refer to resistant, green columns refer to susceptible and yellow column refers to resistant developing *Ae. aegypti*

### Collection of *Aedes* eggs

Eggs were collected using ovitraps baited with a grass infusion. Ovitraps consisted of black 2L containers made of plastic and an oviposition substrate of seed germination paper. The ovitraps were filled with 50ml of 2-3-day old grass infusion and 1200ml of tap or rainwater. After obtaining verbal consents from the household owners, the ovitraps were placed primarily indoors including the main living area (under beds), behind refrigerators, under stairways, in garages, and on balconies. When these sites were not suitable, ovitraps were set in the yards under sheds close to the house. Within Dhaka, the number of ovitraps varied from 50-70 per location, whereas in the areas outside of Dhaka (non-Dhaka), ∼100 ovitraps were set in each location. For all other non-Dhaka districts except Chittagong, eggs were collected from one urban and one rural location. Eggs were collected in 2017 during the traditional peak dengue transmission months from August to November.

### Mosquito rearing

Ovitraps were collected after six days *in situ*. Upon collection, the germination papers were dried and sent to the insectary at the Animal Research Facilities, International Centre for Diarrhoeal Disease Research, Bangladesh (icddr,b) in Dhaka. Due to unexpectedly long time required to prepare the rearing facility, mosquito rearing was delayed until December 2017. Given this delay, hatching rates were low for several locations, so in several cases, eggs from adjacent locations were merged into a single population (Table 1). Mosquitoes were reared at a constant temperature (26-28 °C) and humidity (70-80%). When possible, mosquitoes were reared to the F2 generation to obtain sufficient numbers for a wide range of susceptibility tests. Artificial blood-feeding was provided using the methods described by Costa-da-Silva *et al*. [36]. Adult mosquitoes were provided with 10% sucrose solution. In addition to the field populations, the ‘Rockefeller’ (ROCK) insecticide susceptible *Ae. aegypti* reference strain was obtained from the U.S. Centers for Disease Control and Prevention (CDC, Atlanta, USA) and reared for use as a susceptible control in the bioassays.

**Table 1.** Summary of *Ae. aegypti* populations tested in this study

| Ovitrapping Collection Sites |  |  | Final population*<br>for bioassay | Generation<br>Tested |
| --- | --- | --- | --- | --- |
| Location | District | Location |  |  |
| Dhaka | Dhaka | Azimpur | Azimpur | F0 |
|  |  | Dhanmondi | Dhanmondi &<br>Mohammadpur | F0-F2 |
|  |  | Mohammadpur |  |  |
|  |  | Gulshan | Gulshan & Karail | F0-F2 |
|  |  | Karail |  |  |
|  |  | Mipur | Mipur | F1-F2 |
|  |  | Malibagh | Malibagh | F2 |
|  |  | Uttara | Uttara | F1-F2 |
| Non-Dhaka | Rajshahi | Rajshahi City (Urban) | Rajshahi | F2 |
|  |  | Poba (Rural) |  |  |
|  | Chapai Nawabganj | Chapai Nawabganj City (Urban) | Chapai Nawabganj | F2 |
|  |  | Shibganj (Rural) |  |  |
|  | Bandarban | Bandarban City (Urban) | Bandarban | F0-F2 |
|  |  | Rowangchhari (Rural) | No *Ae. aegypti*, all were *Ae. albopictus* | NA |
|  | Chittagong | Chittagong City | Chittagong | F0-F2 |

\*For ease of description mosquitoes from each location is considered as population

### Insecticide susceptibility testing

Susceptibility tests were conducted following the CDC bottle bioassay protocol [37] using 3-5-day old, non-blood fed female mosquitoes. Four insecticides belonging to three major classes were tested for each population when sufficient mosquitoes were available: the pyrethroids permethrin and deltamethrin, the organophosphate malathion, and the carbamate bendiocarb. Mosquitoes were exposed to the diagnostic dose of the insecticide, and when resistance was detected and sufficient mosquitoes were available, resistance intensity assays were also conducted by exposing mosquitoes to 2X and/or 5X the diagnostic dose. All bioassays comprised >100 mosquitoes per insecticide per population across four test bottles and 15-25 mosquitoes in an untreated control bottle. Susceptibility status was recorded after 0, 15, and 30 minutes of insecticide exposure. Mosquitoes unable to stand were considered dead [37]. Mortality data were interpreted according to World Health Organization (WHO) recommendations, with <90% mortality in a population corresponding to resistance [38].

### Biochemical assays

To detect potential metabolic mechanisms of resistance through the altered activity of detoxifying enzymes, biochemical assays were performed [39]. From each population, 30, 1-2-day old mosquitoes were tested for activities of non-specific β esterase (β-EST), mixed-function oxidases (MFOs), acetylcholine esterase (AChE), and insensitive acetylcholine esterase (IAChE), with a protein assay conducted for each mosquito to control for differences in body size. All mosquitoes were freeze killed and kept at −20 °C until analysis. Briefly, mosquitoes were individually homogenized in 100µl of potassium phosphate buffer followed by dilution to 2ml with additional buffer. For all tests, mosquito homogenates were run in triplicate on 96-well round-bottom microplates (Corning, NY, USA). Homogenates of the Rockefeller (ROCK) susceptible *Ae. aegypti* reference strain was used as a comparator.

For the β-EST assay, 100µl of mosquito homogenate was added in each well followed by 100µl β-naphthyl acetate. The plate was then incubated at room temperature for 20 minutes. After adding 100µl Fast Blue in each well, the plate was further incubated at room temperature for 4 minutes and read by a spectrophotometer (BioTek, VT, USA) using a 540nm filter.

For the MFO assay, 100µl of mosquito homogenate was added to each well followed by 200µl of 3,3,5,5-tetramethylbenzidine (TMBZ) and 25µl 3% hydrogen peroxide. The plate was incubated for 10 minutes and read by a spectrophotometer using a 620nm filter.

For the AChE assay, 100µl of mosquito homogenate was added to each well followed by 100µl of acetylthiocholine iodide (ATCH) and 100µl dithio-bis-2-nitrobenzoic acid (DTNB). The plate was read immediately (T_0_) using a 414nm filter and a second reading was taken after 10 minutes (T_10_). The absorbance at T_0_ was subtracted from T_10_ and used as the value for data analysis.

The IAChE assay was similar to the AChE assay, with the addition of propoxur to the ATCH to quantify the extent to which propoxur inhibited the reaction.

The total protein content of each mosquito was measured by adding 20µl of the homogenate to a well together with 80µl of potassium phosphate and 200µl of protein dye. The plate was read immediately using a 620nm filter.

### DNA extraction

DNA extraction was carried out using the REDExtract-N-Amp™ tissue kit (Merck, Germany) according to the manufacturer’s protocol. Briefly, individual mosquitoes were placed in 1.5ml microcentrifuge tubes and mixed with 100µl of extraction solution and 25µl of tissue preparation solution. Tubes were then incubated at room temperature for 10 minutes followed by further incubation for 3 minutes at 95°C. Then, 100µl of neutralization solution B was added to the sample and the sample was mixed by vortexing.

### Detection of *kdr* alleles (Gly1016, Cys1534, and Leu410)

To understand the correlation between phenotypic resistance and the presence of the *kdr* alleles Gly1016, Cys1534C, and Leu410, phenotyped mosquitoes exposed to permethrin and deltamethrin in the bioassays underwent real-time PCR. An additional 30 non-phenotyped mosquitoes from each of the six Dhaka populations were analyzed to estimate the population-level allele frequencies.

The Gly1016 PCR was performed following the protocol described by Saavedra-Rodriguez *et al*. [40]. Each reaction contained 4.5µl of iQ-SYBR Green Supermix (Bio-Rad Laboratories Inc, CA, USA), 0.45µl of each primer, one common Gly forward (5’-ACC GAC AAA TTG TTT CCC-3’), one reverse primer for either Val (5’-GCG GGC AGC AAG GCT AAG AAA AGG TTA ATT A-3’) or Gly (5’-GCG GGC AGG GCG GGG GCG GGG CCA GCA AGG CTA AGA AAA GGT TAA CTC-3’), 1µl of template DNA and ddH_2_O for a final reaction volume of 9µl. Thermal cycling conditions were: 95°C for 3 min; 40 cycles of 95°C for 10 sec, 58°C for 10 sec, 72°C for 30 sec; 95°C for 10 sec and a ramp from 65°C to 95°C at a rate of 0.2°C/10 sec for melting curve analysis.

The Cys1534 PCR was based on the protocol described by Yanola *et al*. [41]. Each reaction contained 4.5µl of iQ-SYBR Green Supermix (Bio-Rad Laboratories Inc, CA, USA), 0.45µl Cys forward primer (5’-GCG GGC AGG GCG GCG GGG GCG GGG CCT CTA CTT TGT GTT CTT CAT CAT GTG-3’), and 0.45µl each of Phe forward (5′-GCG GGC TCT ACT TTG TGT TCT TCA TCA TAT T-3′) and a common reverse primer (5′-TCT GCT CGT TGA AGT TGT CGA T-3′), 1µl of template DNA and ddH_2_O for a final reaction volume of 9µl. Thermal cycling conditions were: 95°C for 3 min; 40 cycles of 95°C for 10 sec, 57°C for 10 sec, 72°C for 30 sec; 95°C for 10 sec and a ramp from 65°C to 95°C at a rate of 0.5°C/5 sec for melting curve analysis.

The Leu410 PCR was performed based on the protocol described by Saavedra-Rodriguez *et al*. [42]. Each reaction contained 4.5µl of iQ-SYBR Green Supermix (Bio-Rad Laboratories Inc, CA, USA), 0.45µl of each primer, Val forward primer (5’GCG GGC AGG GCG GCG GGG GCG GGG CCA TCT TCT TGG GTT CGT TCT ACC GTG-3’), Leu forward primer (5′-GCG GGC ATC TTC TTG GGT TCG TTC TAC CAT T-3′) and a common reverse primer (5′-TTC TTC CTC GGC GGC CTC TT-3′), 1µl of template DNA and ddH_2_O for a final reaction volume of 9µl. Thermal cycling conditions were: 95°C for 3 min; 40 cycles of 95°C for 10 sec, 60°C for 10 sec, 72°C for 30 sec; 95°C for 10 sec and a ramp from 65°C to 95°C at a rate of 0.2°C/10 sec for melting curve analysis.

### Data analysis

Percent mortality at the diagnostic time of 30 minutes was used to describe the susceptibility status of the mosquito populations tested. Populations were classified as resistant and susceptible based on WHO and CDC guidelines [37, 38]: when mortality was <90% the population was considered as resistant, mortality between 90 - 97% suggested that the population was developing resistance, and mortality ≥98% represented a susceptible population. The 95% confidence intervals (CI) were calculated for the percent mortalities from the bioassays and for allele frequencies.

Interquartile ranges of the mean of the optical density (OD) values from the biochemical assays were compared between the study populations and the susceptible reference strain. Regression analyses were performed to measure the statistical significance of differences between the mean OD values between populations.

Pearson chi-square tests were performed to understand the associations between Gly1016 and Cys1534 genotypes and phenotypes of bioassayed mosquitoes. The population-level allele frequencies were calculated using the following equation [43]:

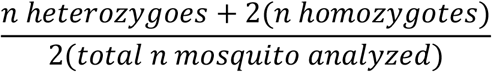

The linkage disequilibrium, departures from the Hardy-Weinberg equilibrium (HWE) and the p-value for Gly1016 and Cys1534 in each population were assessed using Fisher’s exact test in Gene pop (version 4.2) (https://genepop.curtin.edu.au/) [44]. Statistical analyses were conducted in Microsoft Excel 2016 (Microsoft Inc.) and Stata 15 (StataCorp LLC, TX, USA).

## Results

### Insecticide bioassays

In the populations from Dhaka, *Ae. aegypti* mortality ranged between 0% in Malibagh to 6.7% in Gulshan & Karail at the diagnostic dose of permethrin. A higher dose of permethrin (2X the diagnostic dose) was tested with the populations of Dhanmondi & Mohammadpur and Gulshan & Karail but still resulted in <50% mortality at the diagnostic time. In contrast, mortality to deltamethrin varied between areas of Dhaka, ranging from 49.0% (95% CI ± 7.3) in Gulshan & Karail to 100% (95% CI ± 1.6) in Uttara. Susceptibility to malathion was tested in three populations from Dhaka. While the Gulshan & Karail population was resistant (62.9% mortality, 95% CI ± 2.7), the Dhanmondi & Mohammadpur (98.1% mortality, 95% CI ± 2.7) and Uttara (100% mortality, 95% CI ± 2.1) populations were susceptible. All Dhaka populations tested against bendiocarb were susceptible (100% mortality in all populations) (Fig. 1).

The *Ae. aegypti* populations sampled from the non-Dhaka locations were also highly resistant to permethrin, with mortality ranging from 0% in the Chapai Nawabganj population to 14.8% (95% CI ± 2.0) in the Rajshahi population. When the concentration of permethrin was increased to 2X in Chittagong, mortality was still <50%. However, when the permethrin concentration was increased to 5X in Bandarban, the population was fully susceptible (100% mortality). While the Chapai Nawabganj (100% mortality, CI ± 0.79) and Chittagong (99.0% mortality, 95% CI ± 0.79) populations were susceptible to deltamethrin, the Bandarban population was resistant to deltamethrin at the diagnostic dose (67% mortality, 95% CI ± 6.1) but susceptible when the concentration was increased to 2X (99.1% mortality, 95% CI ± 0.79). The Bandarban population was also resistant to the diagnostic dose of malathion (75.7% mortality, 95% CI ± 4.6) but susceptible to malathion 2X (100% mortality). The ROCK strain was confirmed to be fully susceptible to the diagnostic doses of the four insecticides. A summary of bioassay data is presented in Fig. 2.

**Fig. 2.**
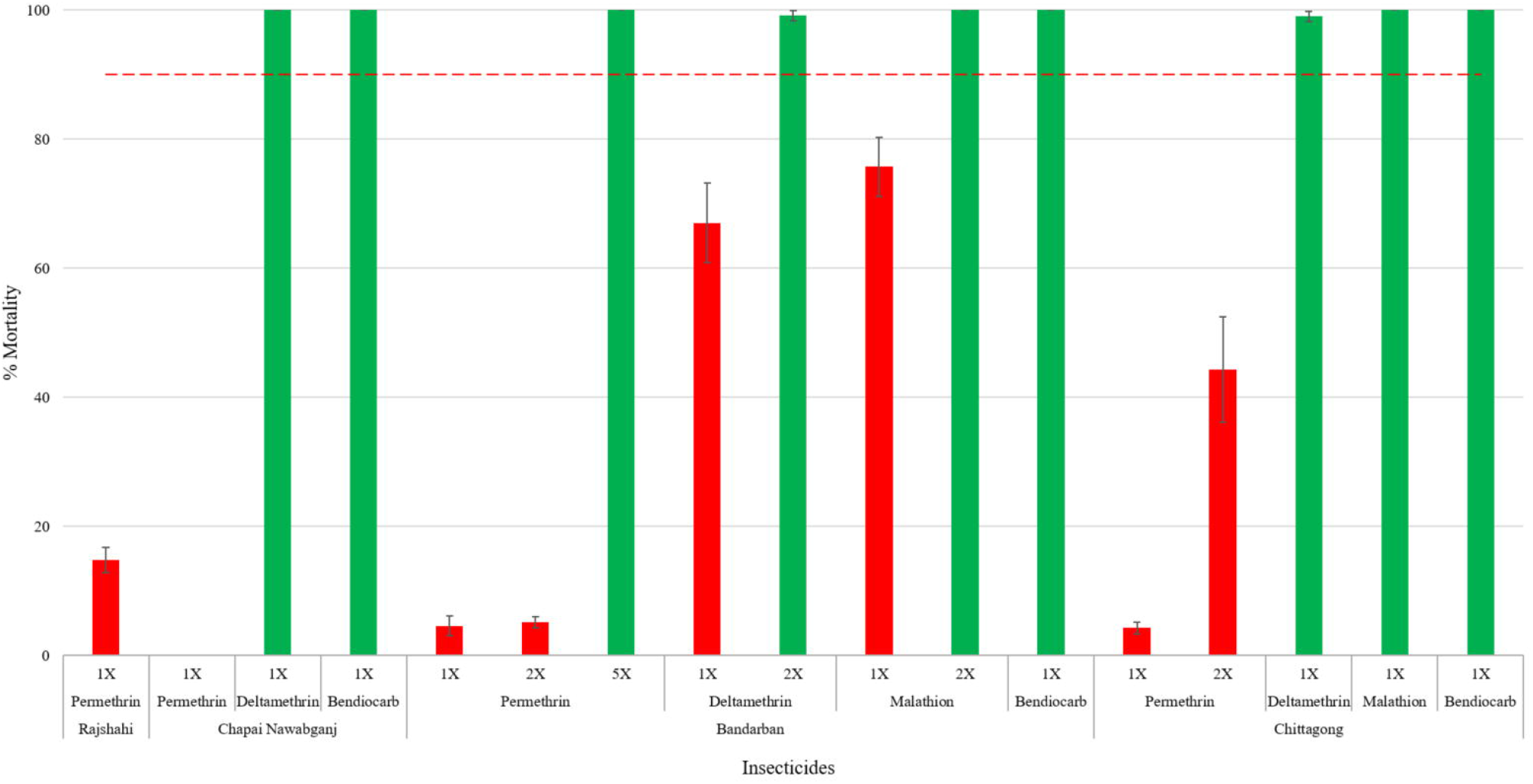
Bioassay results (% Morality at 30 minutes ±95% confidence intervals) for female *Ae. aegypti* from four non-Dhaka locations. Red columns refer to resistant, green columns refer to susceptibile *Ae. aegypti*, red dashed line indicates 90% mortality threshold

### Biochemical assays

All *Ae. aegypti* populations tested from field collections had significantly higher (p<0.0001) MFO activity compared to ROCK. The β-EST activity levels of *Ae. aegypti* populations from Azimpur, Uttara, Dhanmondi & Mohammadpur, Gulshan & Karail, Malibagh, Mirpur, and Bandarban were significantly (p<0.0001) higher than the ROCK reference strain. However, β-EST levels in the non-Dhaka sites of Chapai Nawabganj, Chittagong, and Rajshahi populations were significantly lower than ROCK (p<0.0001). In the case of AChE activity, populations from Azimpur (p<0.042), Chittagong (p<0.019), and Gulshan & Karail (p<0.0001) were significantly higher and Dhanmondi & Mohammadpur (p<0.001) and Malibagh (p<0.001) were significantly lower than ROCK. The estimated levels of IAChE were significantly higher (p<0.0001) in the Gulshan & Karail population compared to ROCK, which suggests that AChE insensitivity may exist in this population. Levels were low across the remaining populations, suggesting that the target site remains sensitive. However, it is noteworthy that levels were significantly lower than ROCK in Bandarban, Chapai Nawabganj, Mirpur, and Uttara (p<0.001). When total protein content was compared between mosquito populations, except Azimpur and Mirpur, all populations were significantly (p<0.026) lower than ROCK, suggesting that the body sizes were generally smaller for most of the field populations (Fig. 3).

**Fig. 3.**
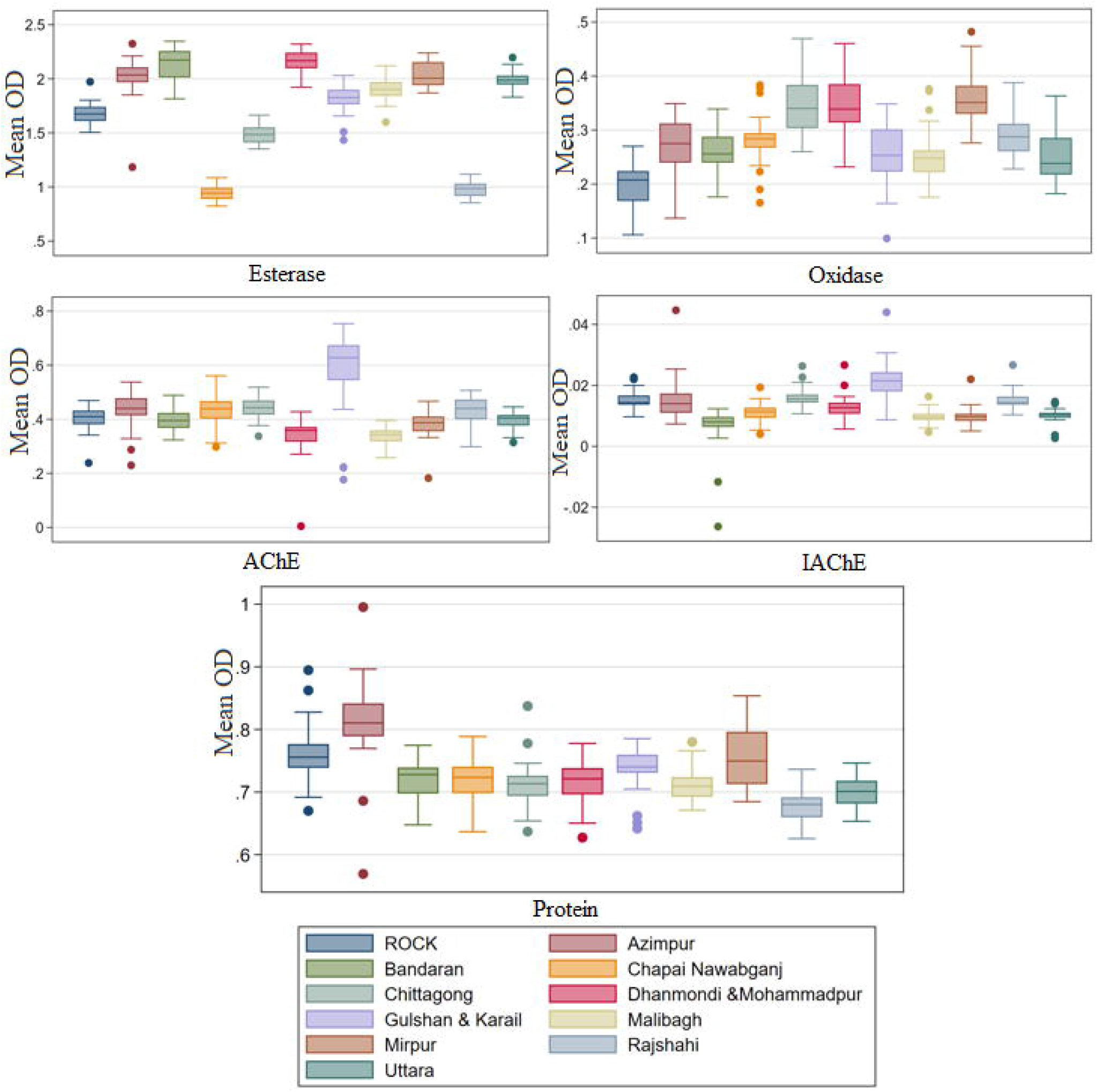
Enzyme activity levels in populations of *Ae. aegypti* from Bangladesh compared to the susceptible ROCK strain. Box plots denote the 50 percentile of the mean OD values, whiskers are the remaining percentile values, and the dots are outliers

### Knockdown resistance (*kdr*) genotyping

A total of 142 phenotyped mosquitoes from permethrin 1X bioassays and 59 phenotyped mosquitoes from deltamethrin 1X bioassays were analyzed for the Gly1016 mutation. From the Dhaka mosquito populations exposed to permethrin, 37.8% (28/74) of the survivors (alive) were mutant homozygotes (GG) and 29.7% (22/74) were wild type homozygotes (VV). The correlations between genotype and phenotype of permethrin-exposed Dhaka mosquitoes were statistically significant (p<0.0001). Most of the dead mosquitoes were wild-type homozygotes (12/14, 85.7%). Amongst the mosquitoes from sites outside of Dhaka, more than half of the permethrin survivors were heterozygotes (23/44, 52.3%,) and there was an equal number (5/10, 50.0%) of wild-type homozygotes and heterozygotes amongst the dead mosquitoes. For deltamethrin, only dead mosquitoes were genotyped due to limitations at the time of the bioassay. Mosquitoes from Dhaka that were dead after exposure to deltamethrin had similar frequencies of all three genotypes. However, the mosquitoes from outside of Dhaka did not include any mutant homozygotes (Table 2).

**Table 2.** Phenotype and genotype at *kdr* locus 1016 in mosquitoes exposed to permethrin and deltamethrin from Dhaka and non-Dhaka populations. GG, mutant homozygotes; VV, wildtype homozygotes; and VG heterozygotes.

|  |  | Permethrin 1X |  | Deltamethrin 1X |
| --- | --- | --- | --- | --- |
| Dhaka | Genotype | Phenotype |  | Phenotype |
|  |  | Alive<br>(n=74) | Dead<br>(n=14) | Dead<br>(n=29) |
|  | VV | 22<br>(29.7 %) | 12<br>(85.7%) | 9<br>(31.0%) |
|  | VG | 24<br>(32.4%) | 1<br>(7.1%) | 9<br>(31.0%) |
|  | GG | 28<br>(37.8%) | 1<br>(7.1%) | 11<br>(37.9%) |
|  | *p* | 0.000 |  |  |
| Non-Dhaka | Genotype | Phenotype |  | Phenotype |
|  |  | Alive<br>(n=44) | Dead<br>(n=10) | Dead<br>(n=30) |
|  | VV | 12<br>(27.3%) | 5<br>(50.0%) | 14<br>(46.7%) |
|  | VG | 23<br>(52.3%) | 5<br>(50.0%) | 16<br>(53.3%) |
|  | GG | 9<br>(20.5%) | 0 | 0 |
|  | *p* | 0.184 |  |  |

Of the 170 mosquitoes screened for Cys1534 mutation, 110 mosquitoes were from permethrin bioassays and the remaining were from deltamethrin bioassays from both Dhaka and non-Dhaka populations. From the permethrin phenotyped Dhaka mosquitoes, 54.1% (33/61) of the resistant (surviving) mosquitoes were 1534 mutant homozygotes (CC) and 41.0% (25/61) were wild type homozygotes (FF). In case of permethrin-susceptible mosquitoes, 90.0% (9/10) were FF and the remaining individual was CC. From the non-Dhaka populations, 37.9% (11/29) of permethrin-resistant mosquitoes were CC and 27.6% (8/29) were heterozygotes (FC). Interestingly, none of the permethrin-susceptible mosquitoes from the non-Dhaka sites was FF, and 8/10 were CC. The correlations between genotype and phenotypes of permethrin exposed mosquitoes in both populations were statistically significant (p<0.016 for Dhaka and p<0.043 for non-Dhaka). A total of 60 dead mosquitoes from the deltamethrin bioassays were analyzed for Cys1534. Interestingly, most of the mosquitoes were wild-type homozygotes FF (65.5%, 19/30) in Dhaka, whereas, the opposite was seen for non-Dhaka populations (Table 3).

**Table 3.** Phenotype and genotype at *kdr* locus 1534 in mosquitoes exposed to permethrin and deltamethrin from Dhaka and non-Dhaka populations. CC, mutant homozygotes; FF, wildtype homozygotes; and FC heterozygotes.

|  |  | Permethrin 1X |  | Deltamethrin 1X |
| --- | --- | --- | --- | --- |
| Dhaka | Genotype | Phenotype |  | Phenotype |
|  |  | Alive<br>(n=61) | Dead<br>(n=10) | Dead<br>(n=30) |
|  | FF | 25<br>(41.0%) | 9<br>(90.0%) | 19<br>(63.3%) |
|  | FC | 3<br>(4.9%) | 0 | 0 |
|  | CC | 33<br>(54.1%) | 1<br>(10.0%) | 11<br>(36.7%) |
|  | *p* | 0.016 |  |  |
| Non-Dhaka | Genotype | Phenotype |  | Phenotype |
|  |  | Alive<br>(n=29) | Dead<br>(n=10) | Dead<br>(n=30) |
|  | FF | 10<br>(34.5%) | 0 | 2<br>(6.7%) |
|  | FC | 8<br>(27.6%) | 2<br>(20.0%) | 9<br>(30.0%) |
|  | CC | 11<br>(37.9%) | 8<br>(80.0%) | 19<br>(63.3%) |
|  | *p* | 0.043 |  |  |

All mosquitoes (n=264) from permethrin and deltamethrin bioassays (1X and 2X) genotyped for Leu410 were found to be wild-type homozygotes.

Of the 177 non-phenotyped mosquitoes from the Dhaka populations, more than half were V1016G heterozygotes (51%, n=90/177). The highest Gly1016 homozygote (GG) frequency was observed in Gulshan & Karail (77%, n=23/30) followed by Mirpur (47%, n=14/30) and Malibagh (38%, n=11/29) (Fig. 4). In the case of Cys1534, the largest group of the mosquitoes were homozygous wild type (FF) (43.5%, n=77/177). The highest mutant homozygote (CC) frequency was recorded from Dhanmondi & Mohammadpur (41.4%, n=12/29) (Fig. 5).

**Fig. 4.**
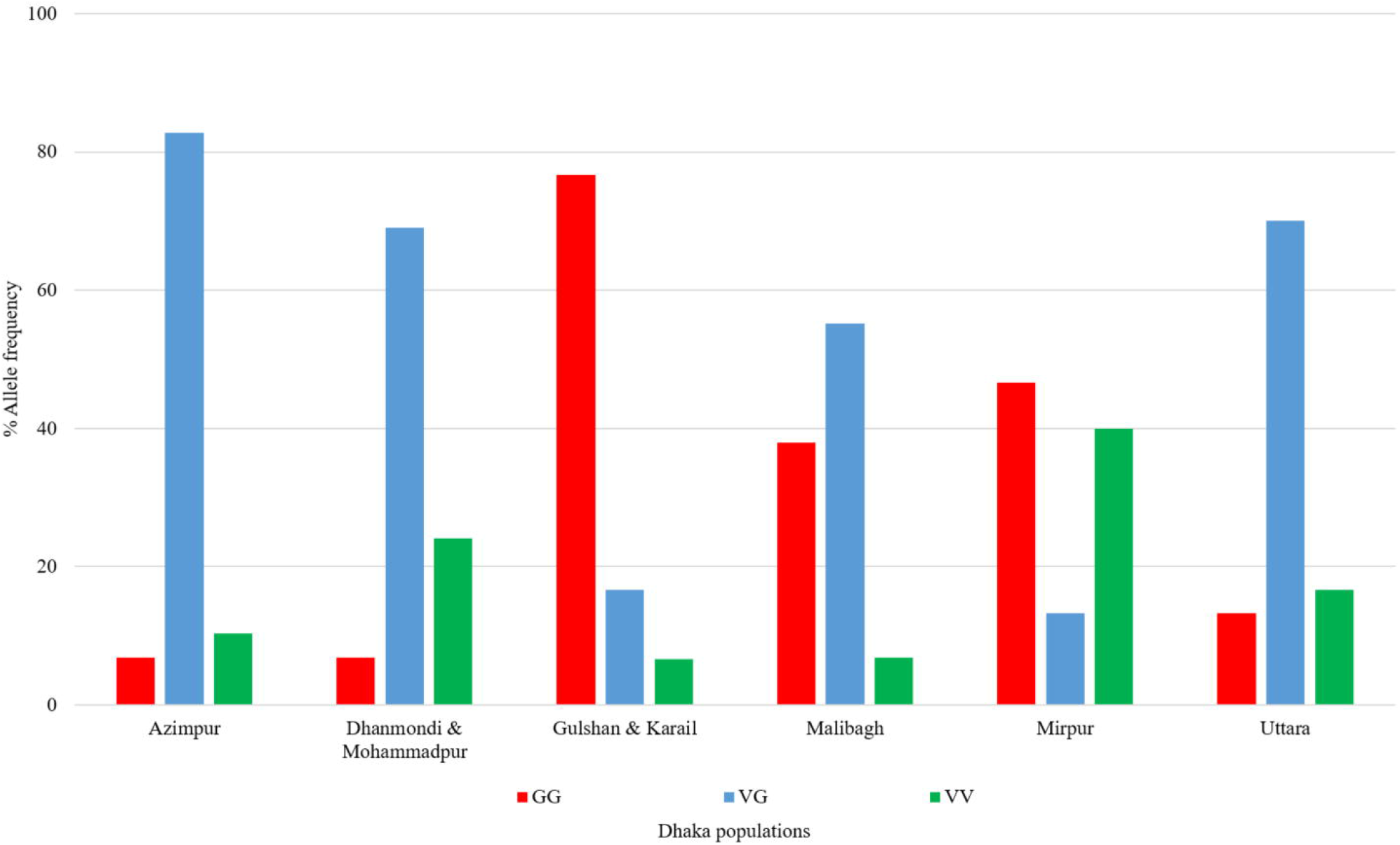
Allele frequencies of Gly1016 in *Ae. aegypti* populations from Dhaka

**Fig. 5.**
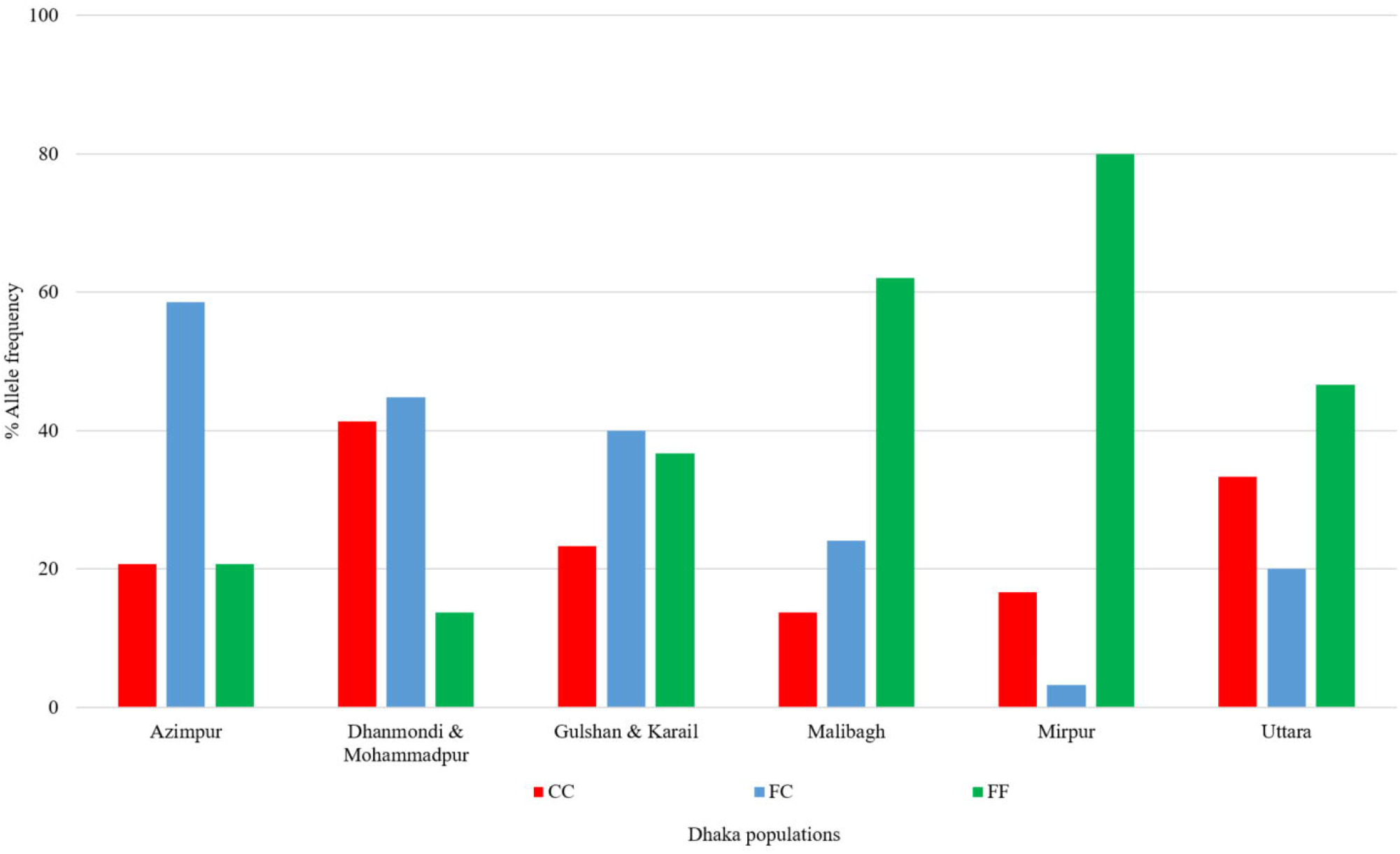
Allele frequencies of Cys1534 in *Ae. aegypti* populations from Dhaka

The overall allele frequency of Gly1016 was 57.1% (95% CI ± 8.41) and of Cys1534 was 38.4% (95% CI ± 5.66). The highest frequency of Gly1016 was 85.0% (95% CI ± 30.42) in Gulshan & Karail. The highest frequency of Cys1534 was 63.8% (95% CI ± 23.22) in Dhanmondi & Mohammadpur (Table 4). The Hardy-Weinberg equilibrium (HWE) test revealed that three populations had significant departures from HWE. This includes the Azimpur population for Gly1016, the Uttara population for Cys1534, and the Mirpur population for both (Table 4).

**Table 4.** Frequency of Gly1016 (GG) and Cys1534 (CC) *kdr* alleles in *Ae. aegypti* populations from Dhaka

| Populations | Allele | n | % Frequency | 95% CI | p-value of HWE |
| --- | --- | --- | --- | --- | --- |
| Azimpur | G | 29 | 48.3 | 17.6 | 0.0008* |
|  | C |  | 50.0 | 18.2 | 0.4719 |
| Dhanmondi & Mohammadpur | G | 29 | 41.4 | 15.1 | 0.0526 |
|  | C |  | 63.8 | 23.22 | 1.000 |
| Gulshan & Karail | G | 30 | 85.0 | 30.4 | 0.0991 |
|  | C |  | 43.3 | 15.5 | 0.4540 |
| Malibagh | G | 29 | 65.5 | 23.9 | 0.4194 |
|  | C |  | 25.9 | 9.4 | 0.0546 |
| Mirpur | G | 30 | 55.3 | 19.1 | 0.0000* |
|  | C |  | 18.3 | 6.6 | 0.0001* |
| Uttara | G | 30 | 48.3 | 17.3 | 0.0654 |
|  | C |  | 43.3 | 15.5 | 0.0024* |

\* Significant p-values

## Discussion

The application of chemical insecticides either in the form of space sprays, thermal fogging, or LLINs has been carried out for many years in Bangladesh. However, documents of mosquito susceptibility to insecticides are scanty. Some information can be obtained from the ‘Malaria Threat Map’ website about insecticide resistance in some *Anopheles* species [45]. A recent article reported permethrin and deltamethrin resistance in the highest malarious region in *Anopheles vagus* [34]. However, these reports are limited to phenotypic characteristics and no clear understanding of resistance mechanisms for any mosquito species is available. Despite the increasing prevalence of *Aedes*-borne diseases in Bangladesh, the insecticide resistance status of *Ae. aegypti* has previously not been assessed. The results reported here provide a comprehensive overview of insecticide resistance across Dhaka, and in several other sites of high epidemiological importance. We report a high frequency and intensity of permethrin resistance in all populations that were studied. Despite this high level of permethrin resistance, susceptibility to deltamethrin was still present in several of the populations. This difference suggests that the underlying mechanisms causing resistant phenotypes in these populations may not be shared across the pyrethroid class.

The increased activity of enzymes including β esterases and mixed-function oxidases in the populations suggests an important role of metabolic mechanisms in conferring resistance. All Dhaka populations had elevated levels of esterase and oxidase activity and were resistant to permethrin. Outside of Dhaka, esterase activity was notably lower in Chapai Nawabganj and Rajshahi, and while both populations were resistant to permethrin, the latter population remained susceptible to deltamethrin (the former was not tested). Increased activity of esterases and oxidases may also be associated with the malathion resistance that was detected in Gulshan & Karail and Bandarban [46, 47, 48, 49]. In addition, AChE activity was elevated in Gulshan & Karail and could also be contributing to the malathion resistance that was detected there. An important limitation of the biochemical assay data is the lack of information on glutathione S-transferases (GSTs). A growing body of evidence suggests that these are important mechanisms in pyrethroid resistance in *Ae. aegypti* [50], with *GSTe2* associated with resistance to both permethrin and deltamethrin, and *GSTe7* with deltamethrin [51, 52].

The Gly1016 and Cys1534 *kdr* mutations have been widely reported in Asia [41, 53, 54]. An additional mutation, Leu410, has also been reported in association with pyrethroid resistance, but its prevalence in Asia has not yet been well studied [23]. Expression of insect sodium channels in *Xenopus* oocytes coupled with electro-neurophysiological measurements has demonstrated that Gly1016, Cys1534, and Leu410 reduce the sensitivity of the VGSC to permethrin and deltamethrin [23, 55]. However, Leu410 was not detected in any of the populations in the current study. This is unexpected, as previous research has suggested the parallel evolution of this mutation together with the polymorphisms at positions 1016 and 1534 [42], both of which were detected at moderate to high frequencies in our study. The co-occurrence of Pro989 with Gly1016 conferring high pyrethroid resistance in *Ae. aegypti* has been reported previously [56]. However, this current study did not include S989P *kdr* detection.

The *kdr* mutations Gly1016 and Cys1534 were found at varying frequencies across Dhaka. This fine-scale spatial heterogeneity suggests that selection pressures for insecticide resistance are variable across small spatial scales within Dhaka, and reflects trends that have been reported elsewhere [43, 57]. Historically, *Aedes* control in Dhaka and major cities in Bangladesh solely depends on thermal fogging using a combination of pyrethroid insecticides. Pyrethroids are also commonly used in households via commercially available coils and aerosols. Both operational and domestic insecticide use may contribute to insecticide resistance selection pressures in *Ae. aegypti* [58].

In Dhaka, 1016G and 1534C homozygous mutants were mostly associated with survival in the permethrin bioassays. It is also worth noting that the population from the Dhaka neighborhood of Mirpur was resistant to permethrin yet susceptible to deltamethrin and was also the population with the highest frequency of Val1016 and Phe1534 wild-type homozygotes. These findings suggest that while *kdr* alleles may be contributing to the insecticide resistance that was detected, they are not the only mechanism and such relationship is not rare [56, 59].

From an operational perspective, the data presented here will be important in guiding the choice of vector control tools. Given the widespread and intense permethrin resistance that was detected, vector control products containing alternative compounds should be used. Although some populations remained susceptible to deltamethrin, given the high degree of permethrin resistance, it would be prudent to search for alternatives outside of the pyrethroid class. Particularly notable was the detection of deltamethrin resistance in Bandarban, where deltamethrin-treated bed nets are routinely used for malaria control [34]. Bandarban was also the only non-Dhaka site to show significantly elevated esterase activity, suggesting that the population was experiencing comparatively greater selective pressure across multiple mechanisms as compared to the other non-Dhaka sites. Vector control activities have focused largely on malaria vectors and have not routinely targeted *Aedes* in this part of Bangladesh. The finding that the *Aedes* population was resistant to the insecticide relied upon for malaria control highlights the importance of implementing strategies based on integrated vector management in Bandarban.

The only insecticide to which every population tested was susceptible was bendiocarb. However, there is no product registered in Bangladesh that could be employed for *Aedes* control that contains bendiocarb as an active ingredient. Therefore, malathion came out as the next best candidate, as public health agencies were desperately seeking alternatives to pyrethroids. Nevertheless, malathion resistance was detected in several of the populations studied, both inside and outside of Dhaka. Also, malathion has been used in agriculture for many years in Bangladesh, so selection pressure outside of vector control already exists to a certain degree [60, 61]. In such a scenario as we detected in Bangladesh with a patchwork of insecticide-resistant phenotypes, it will be challenging to find a ‘one size fits all’ solution for *Aedes* control.

## Conclusion

This current study provides evidence of insecticide resistance in *Ae. aegypti* and data on resistance mechanisms including detoxification enzymes and *kdr* mutations in Bangladesh. High pyrethroid resistance may be compromising the existing *Aedes* control strategies, and the presence of multiple resistance mechanisms poses further challenges regarding alternatives. Continuous surveillance of insecticide resistance will enable trends in susceptibility to be monitored over space and time and will provide a more robust evidence base upon which to select the most effective vector control tools and strategies. In cities like Dhaka where operational control faces challenges posed by insecticide resistance, in addition to the rational use of chemicals, sustainable and alternative tools like biocontrol approaches should be considered.

### Impact

The preliminary results were disseminated among different stakeholders and mosquito control authorities immediately after analyzing the data. Followed by the outbreak of dengue during the monsoon season of 2019 this research findings and recommendations were reinvestigated by the policymakers. As a result, permethrin was replaced by malathion for the control of adult mosquitoes in Dhaka city [62, 63].

## Abbreviations

AChE: acetylcholine esterase;
BI: Breteau Index;
β-EST: β esterase;
CI: confidence intervals;
DDT: dichlorodiphenyltrichloroethane;
DTNB: dithio-bis-2-nitrobenzoic acid;
GSTs: glutathione S-transferases;
HWE: Hardy-Weinberg equilibrium;
IRS: indoor residual spraying;
IACHE: insensitive acetylcholine esterase;
icddr,b: International Centre for Diarrhoeal Disease Research, Bangladesh;
kdr: knockdown resistance:
LLINs: long-lasting insecticidal nets:
MFOs: mixed-function oxidases;
OD: optical density;
ROCK: Rockefeller;
CDC: U.S. Centers for Disease Control and Prevention;
VGSC: voltage-gated sodium channel;
WHO: World Health Organization

## Acknowledgments

icddr,b acknowledges with gratitude the commitment of CDC to its research efforts. icddr,b is also grateful to the Governments of Bangladesh, Canada, Sweden, and the UK for providing core/unrestricted support.

## Authors’ Contributions

HMA, SI, MSA, and AL conceived and designed the study. HMA, FTJ, LV and MRHH carried out field and laboratory work. HMA, MSA and AL coordinated the study. KKP, WAK and RH supervised the study. HMA, MSA, and AL analysed and drafted the manuscript. All authors reviewed the manuscript to its current form. All authords read and approved the final manuscript.

## Availability of data and materials

All data generated or analyzed during this study are included in this published article.

## Ethics approval and consent to participate

The study was approved by the icddr,b Ethical Review Committee (ERC), PR-17050 and informed oral consent was obtained from the household owners before ovitrap installation.

## Consent for publication

Not applicable.

## Disclaimer

The findings and conclusions in this paper are those of the authors and do not necessarily represent the official position of the CDC.

